## Supplementary materials for "Hasty sensorimotor decisions rely on an overlap of broad and selective changes in motor activity"

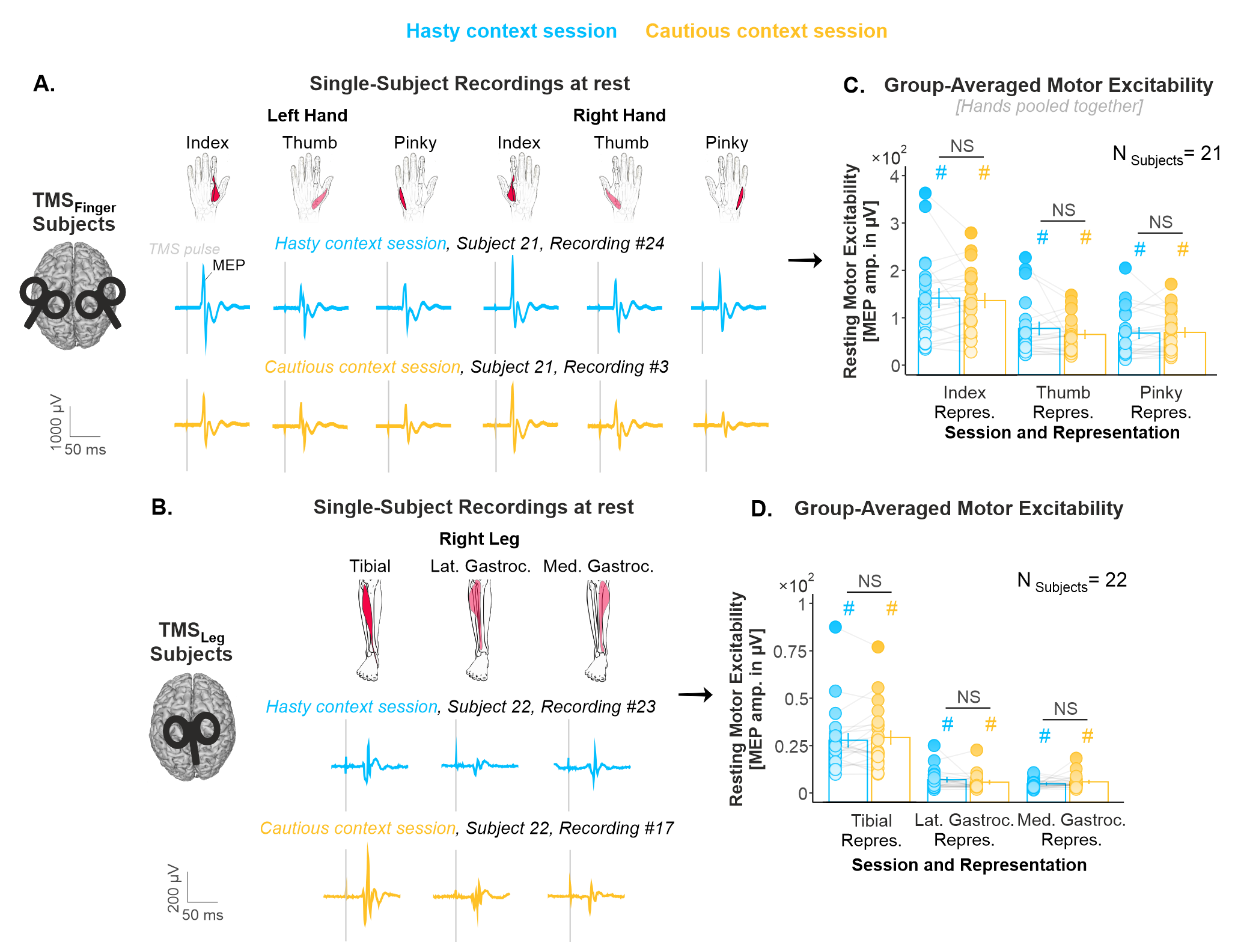

**S1 Figure (related to Figure 1.C): Resting-state MEPs. A. Resting-state recordings in a representative TMS_Finger_ subject.** As described in the Methods section, subjects performed the two block types (*i.e.*, hasty and cautious context blocks) on separate experimental sessions. The blue and yellow traces represent raw MEP recordings, as obtained at rest in the hasty and cautious context sessions, respectively. In each session, the double-coil stimulation over the left and right finger representations allowed us to elicit MEPs in the index, the thumb and the pinky muscles of both hands at once. **B. Same as A for a TMS_Leg_ subject.** MEP amplitudes were smaller in the leg than in the finger representations, potentially due to the higher distance between the coil and the leg area, located in the interhemispheric fissure. Still, in each session, the stimulation over the left leg representations allowed us to elicit MEPs of reliable amplitudes in the right tibialis anterior, as well as in the right lateral and medial heads of the gastrocnemius muscle at once. **C and D.** **Group-averaged resting-state excitability and statistical analysis.** NS annotations indicate that the repeated-measures [rm]ANOVAs performed on resting-state MEPs did not show any significant difference between the hasty and cautious sessions, neither in TMS_Finger_ subjects (Effect of SESSION: F_1,20_ = 0.48, p = .497, partial η^2^ = .023; SESSION*REPRESENTATION interaction: F_2,40_ = 1.08, p = .348, partial η^2^ = .051), nor in TMS_Leg_ subjects (Effect of SESSION: F_1,21_ = 0.19, p = .663, partial η^2^ = .009; SESSION*REPRESENTATION interaction: F_2,42_ = 2.07, p = .138, partial η^2^ = .089). Further, a Bayes Factor (BF) analysis provided substantial evidence for a lack of effect of the factor SESSION on resting-state MEPs (BFs = 5.58 and 5.30, in TMS_Finger_ and TMS_Leg_ subjects, respectively). The hash signs above the bars indicate that MEP amplitudes were significantly higher than 0 in all muscles (all t-values > 5.5, all p-values < .0001 after Bonferroni correction). Error bars represent 1 SEM. All individual and group-averaged numerical data exploited for S1 Figure are freely available at this link <https://osf.io/tbw7h/> (‘Figure_S1_Data.xlsx’).

Altogether, these data show that the two TMS protocols allowed us to record MEPs that were both reproducible across sessions and of reliable amplitudes in all of the investigated muscles.

| **Interaction tested** | **Key statistics** | **DT** | **Accuracy** | | **Urgency Intercept** |
| --- | --- | --- | --- | --- | --- |
| **CONTEXT * SESSION-ORDER** | F-value | 0.0004 | 1.30 | 2.82 | |
|  | p-value | .984 | .259 | .099 | |
|  | **Bayes Factor** | **3.48** | **4.45** | **3.22** | |

**S2 Table (related to Figure 2): The context-dependent shift in SAT did not depend on the session.** As mentioned in the Results section, the session order was not completely counterbalanced among the 50 subjects included in the behavioral analysis: 24 subjects started the experiment with the hasty session while 26 started with the cautious one. To ensure that the effects of CONTEXT observed on DT, accuracy, urgency intercept reported in Figure 2 did not depend on the lack of counterbalancing, we performed Bayesian rmANOVAs, testing whether any of these effects interacted with the factor SESSION-ORDER. We did not find any significant CONTEXT*SESSION-ORDER interaction, whether looking at DTs (F_1, 48_ = 0.00043, p = .984, partial η^2^ = 8.81×10^-5^), at accuracy (F_1, 48_ = 1.30, p = .259, partial η^2^ = .026), or at urgency intercepts (F_1, 48_ = 1.30, p = .259, partial η^2^ = .055). Bayes Factors for these three variables were 3.48, 4.45 and 3.22, providing strong evidence for a lack of CONTEXT*SESSION-ORDER interaction on these variables.

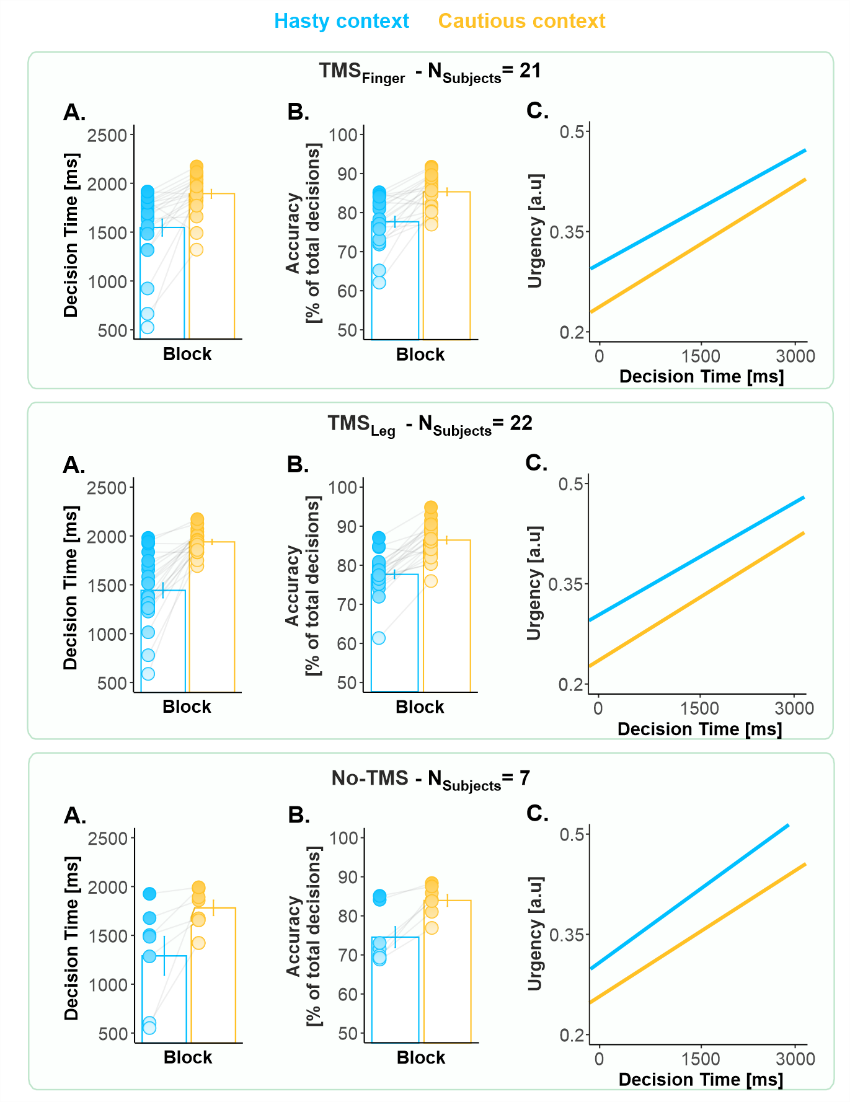

**S3 Figure (related to Figure 2)**: **The context-dependent shift in decision behavior was comparable in the three TMS subgroups (*i.e.*, TMS_Finger_ [top panel], TMS_Leg_ [middle panel] and No-TMS subjects [bottom panel]).** **A. Decision times.** We performed a rmANOVA while considering TMS-SUBGROUP as a categorical predictor. We did not find any significant effect of TMS-SUBGROUP (F_2,47_ = 1.39, p = .258, partial η^2^ = .056) nor of its interaction with the factor CONTEXT on DTs (F_2,47_ = 1.03, p = .362, partial η^2^ = .042). Further, a BF analysis provided substantial evidence for a lack of effect of the TMS-SUBGROUP on DTs (BF = 4.06). **B. Same as A. for decision accuracy.** There was no significant effect of TMS-SUBGROUP (F_2,47_ = 1.05, p = .357, partial η^2^ = .043) nor of its interaction with the factor CONTEXT on accuracy (F_2,47_ = 0.38, p = .687, partial η^2^ = .016). The BF was of 4.07 for the effect of the TMS-SUBGROUP, revealing substantial evidence for a lack of effect of this factor on accuracy. **C. Urgency functions.** There was also no significant effect of TMS-SUBGROUP (F_2,47_ = 0.89, p = .415, partial η^2^ = .037) nor of its interaction with the factor CONTEXT on the slope of the urgency functions (F_2,47_ = 0.81, p = .452, partial η^2^ = .033). Similarly, there was no significant effect of TMS-SUBGROUP (F_2,47_ = 0.19, p = .820, partial η^2^ = .008) nor of its interaction with the factor CONTEXT on the intercept of the functions (F_2,47_ = 0.44, p = .643, partial η^2^ = .018). Here again, BFs showed substantial evidence for a lack of effect of the TMS-SUBGROUP on the slope and the intercept of the functions (BFs = 4.22 and 8.35, respectively). Error bars represent 1 SEM. All individual and group-averaged numerical data exploited for S3 Figure are freely available at this link <https://osf.io/tbw7h/> (‘Figure_S3_Data.xlsx’).

Overall, these results highlight that the three subgroups presented very similar effects of context on all of these behavioral variables, indicating that the application of TMS over the finger and leg representations did not perturb SAT regulation in our task.

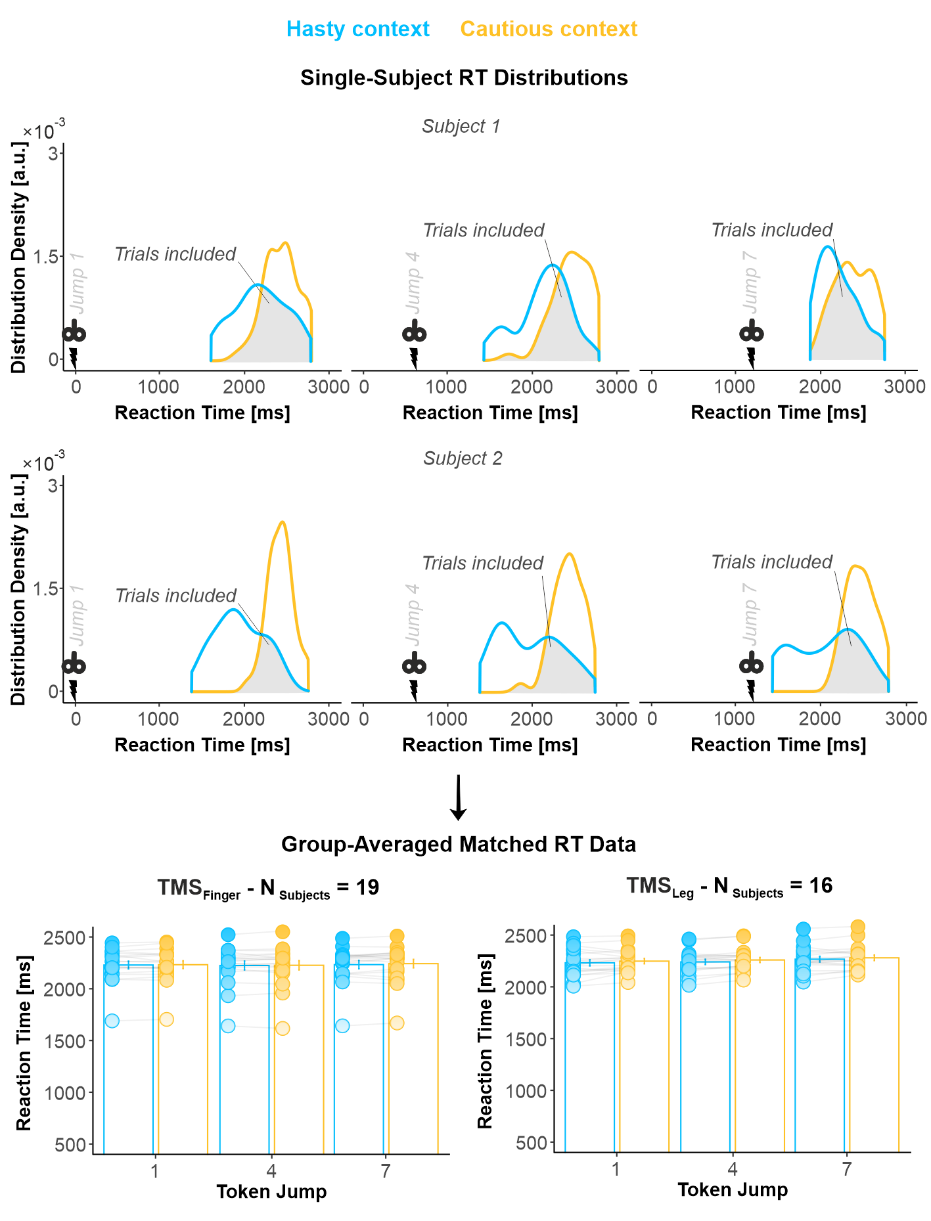

**S4 Figure (related to Figure 3)**: **Trial selection and RT-matching procedure for MEP analysis.** Given the between-context difference in decision speed (see Figure 2 and S2) and its potential effect on MEP amplitudes, we adopted a RT-matching procedure to homogenize RT distributions across contexts (see Murphy et al., 2016, Nat. Comm., for a similar procedure). The procedure consisted in discretizing each subject’s RT distributions into bins of 200 ms width and, for each bin, randomly selecting a matched number of trials. To do so, for each bin, we kept all the trials of the context condition that had the lowest trial count and selected a matched number from the context condition that had the greatest trial count in that bin. As a result, the MEPs included in the analysis were those for which the RT distributions for the hasty and cautious overlapped (grey area on single-subject distributions). In a few subjects (for whom the distribution overlap was very small), this procedure led to the exclusion of many trials. Hence, we had to exclude 2 and 6 subjects out of the 21 TMS_Finger_ and 22 TMS_Leg_ participants as they presented too few trials after this procedure for each timing and context (*i.e.*, < 8 trials on average). On the remaining 19 TMS_Finger_ and 16 TMS_Leg_ subjects, the included trials involved comparable RTs in the hasty and cautious contexts, both in TMS_Finger_ subjects (2230 ± 39 ms and 2230 ± 38 ms, respectively) and in TMS_Leg_ subjects (2266 ± 27 ms and 2278 ± 26 ms, respectively; see bar graphs). Error bars represent 1 SEM. All individual and group-averaged numerical data exploited for S4 Figure are freely available at this link <https://osf.io/tbw7h/> (‘Figure_S4_Data.xlsx’).

Hence, this procedure guaranteed that any effect of context on MEP amplitudes could not result from a between-context difference in RT in the included trials.

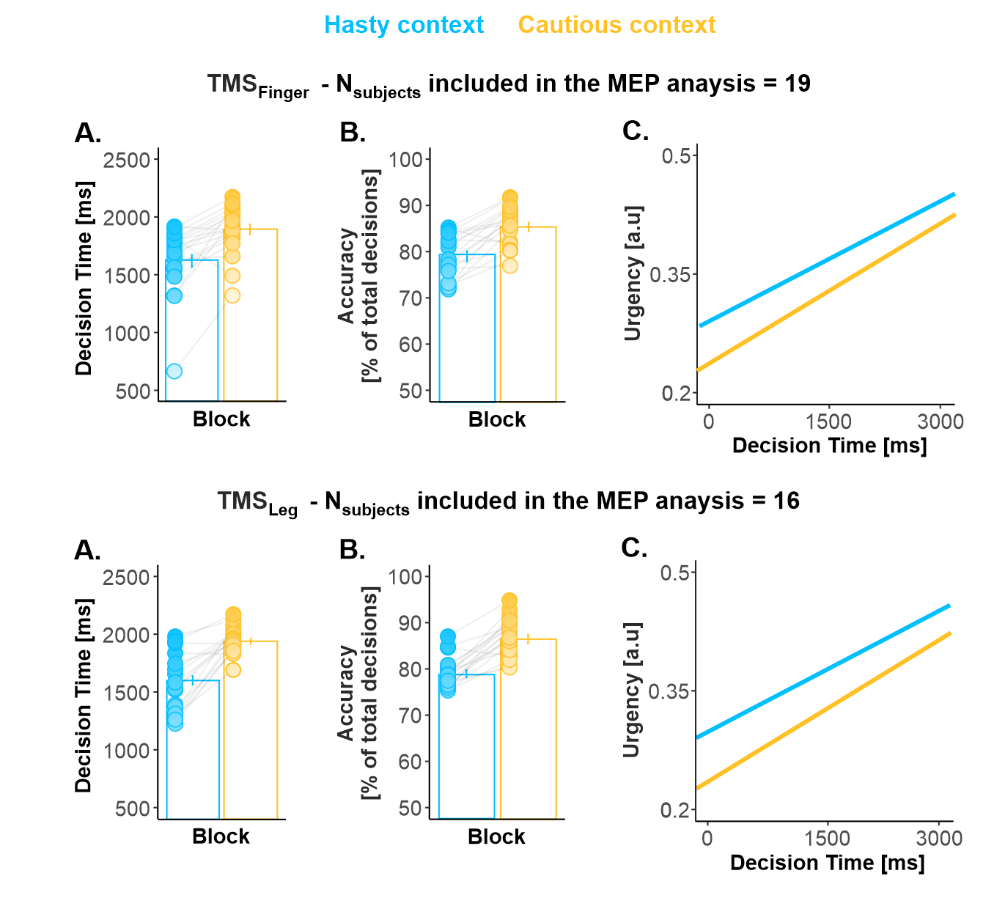

**S5 Figure (related to Figure 4)**: **The context-dependent shift in decision behavior was present in the subjects included in the MEP analysis.** As mentioned in the Results and Methods sections, we had to exclude 3 out of the 21 TMS_Finger_ subjects and 6 out of the 22 TMS_Leg_ subjects following the RT-matching procedure. Subjects excluded following this procedure were more likely to present a too small overlap between their RT distributions and thus to exhibit strong SAT shifts. To ensure that the subjects included in the MEP analysis presented a SAT shift, we performed a statistical analysis on their behavioral data. A between-context comparison revealed that DTs and accuracy were significantly lower in the hasty context (TMS_Finger_ and TMS_Leg_ subjects pooled together: t_34_ = -7.69, p < .0001, Cohen’s d = 1.304 and t_34_ = -9.25, p < .0001, Cohen’s d = 1.565, respectively; panel A and B). Besides, the urgency intercept was significantly higher in the hasty relative to the cautious context (t_49_ = 5.37, p < .0001, Cohen’s d = 0.909; panel C). All individual and group-averaged numerical data exploited for S5 Figure are freely available at this link <https://osf.io/tbw7h/> (‘Figure_S5_Data.xlsx’).

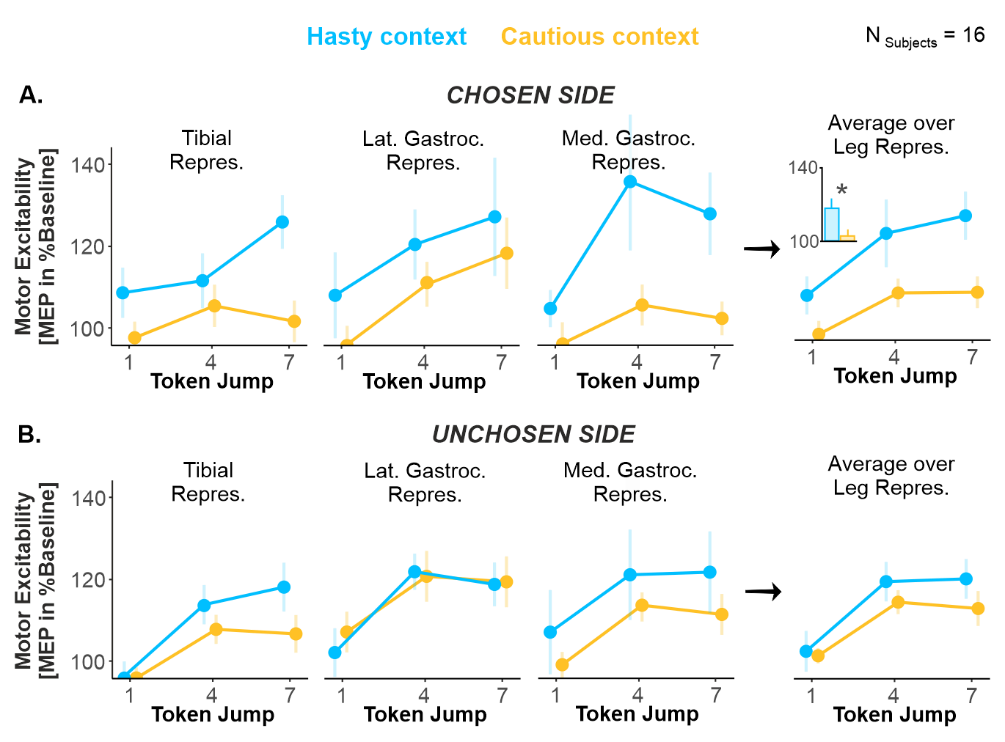

**S6 Figure (related to Figure 4)**: **The broad amplification affected the three leg representations of the chosen side in a reproducible way. A. Effect of CONTEXT on motor excitability on the chosen side.** Excitability changes were more variable in the leg than in the finger representations (*i.e.*, compared to Figure 4), potentially due to the smaller MEP amplitudes obtained for the leg representation (see Figure S1). Despite this variability, the effect of context was comparable in the three investigated leg representations, with higher excitability values in the hasty than in the cautious context. As such, there was no significant CONTEXT*REPRESENTATION interaction (F_2,30_ = 0.53, p = .595, partial η^2^ = .034), nor any CONTEXT*TIMING*REPRESENTATION interaction (F_4,60_ = 1.53, p = .202, partial η^2^ = .093); BF for the latter analysis was of 14.03, providing strong evidence for a lack of effect on this interaction. *: significant effect of context at p < .05. **B. Same as A. for the unchosen side.** Here again, the three leg representations exhibited similar patterns of excitability changes, with no evident impact of context and an overall rise as time elapsed. Indeed, there was no significant CONTEXT*REPRESENTATION interaction (F_2,30_ = 0.58, p = .561, partial η^2^ = .037), nor any CONTEXT*TIMING*REPRESENTATION interaction (F_4,60_ = 0.47, p = .756, partial η^2^ = .030); BF for the latter analysis was of 17.65, providing strong evidence for a lack of effect on this interaction. Error bars represent 1 SEM. All individual and group-averaged numerical data exploited for S6 Figure are freely available at this link <https://osf.io/tbw7h/> (‘Figure_S6_Data.xlsx’).

| **Effect tested** | **Key statistics** | **Motor excitability on the chosen side** | | |
| --- | --- | --- | --- | --- |
|  |  | **Index** | **Thumb** | **Pinky** |
| **CONTEXT** | F-value | 1.17 | **4.34** | 1.82 |
|  | p-value | .294 | **.051** | .194 |
| **TIMING** | F-value | **13.16** | **3.38** | **6.91** |
|  | p-value | **.00005** | **.045** | **.003** |
| **CONTEXT * TIMING** | F-value | **3.94** | 0.44 | 0.62 |
|  | p-value | **.028** | .646 | .543 |

**S7 Table (related to Figure 4): Decomposition of the CONTEXT*TIMING*REPRESENTATION interaction using separate rmANOVAs for each representation with CONTEXT and TIMING as within-subject factors.** For the sake of homogeneity, the same procedure was applied throughout the manuscript to decompose significant effects following interactions. That is, Tukey HSD post-hoc tests were applied for all pairs of conditions comprised in the interaction, and the correction was thus proportional to the number of pairs tested. Here, we exploited an alternative decomposition of the CONTEXT*TIMING*REPRESENTATION interaction found on the chosen side of TMS_Finger_ subjects, using three separate rmANOVAs for each representation with CONTEXT and TIMING as within-subject factors. First, this analysis showed that the global effect of TIMING reported in the manuscript (Figure 3) could be replicated for the three representations using this approach. Further, a CONTEXT*TIMING interaction was present in the index representation (F_2,36_ = 3.94, p = .028, partial η^2^ = .181), consistent with the effects reported in the manuscript (Figure 4.A, left panel). However, this interaction was not present for the surrounding finger representations, neither for the thumb (F_2,36_ = 0.44, p = .646, partial η^2^ = .024) nor for the pinky one (F_2,36_ = 0.624, p = .543, partial η^2^ = .033). Interestingly, the thumb representation presented a main effect of CONTEXT (marginally significant: F_1,18_ = 4.34, p = .051, partial η^2^ = .194), while this effect was not significant for the pinky representation (F_1,18_ = 1.82, p = .194, partial η^2^ = .089). Altogether, this analysis may suggest that the surround suppression effect reported in the manuscript was the strongest in the thumb representation, putatively due to its stronger functional link with the index representation. The analysis also hints that, in the thumb representation, the suppression of excitability did not necessarily depend on the timing at which it was probed and was potentially already present early on during the decision process (*i.e.*, see Jump_1_ in Figure 4.A).

| **Interaction tested** | **Key statistics** | **Motor Excitability on the chosen side** |
| --- | --- | --- |
| **CONTEXT * SESSION-ORDER** | F-value | 2.16 |
|  | p-value | .159 |
|  | **Bayes Factor** | **3.08** |
| **CONTEXT * REPRESENTATION * SESSION-ORDER** | F-value | 1.48 |
|  | p-value | .240 |
|  | **Bayes Factor** | **30.09** |
| **CONTEXT * TIMING * SESSION-ORDER** | F-value | 2.49 |
|  | p-value | .097 |
|  | **Bayes Factor** | **8.63** |
| **CONTEXT * REPRESENTATION * TIMING * SESSION-ORDER** | F-value | 0.32 |
|  | p-value | .863 |
|  | **Bayes Factor** | **10.09** |

**S8 Table (related to Figure 4): In TMS_Finger_ subjects, the effect of context on motor excitability observed on the chosen side did not depend on the session order.** As mentioned in the Results section, the session order was not completely counterbalanced among the 19 TMS_Finger_ subjects included in the MEP analysis: 8 subjects started the experiment with the hasty session while 11 started with the cautious one. To ensure that the effects of CONTEXT observed on motor excitability in these subjects did not depend on the lack of counterbalancing, we performed Bayesian rmANOVAs, testing whether the factor CONTEXT interacted with SESSION-ORDER. We did not find any significant CONTEXT*SESSION-ORDER (F_1, 17_ = 2.16, p = .159, partial η^2^ = .113), CONTEXT*REPRESENTATION *SESSION-ORDER (F_2, 34_ = 1.48, p = .240, partial η^2^ = .081), CONTEXT*TIMING *SESSION-ORDER (F_2, 34_ = 2.49, p = .097, partial η^2^ = .128), or CONTEXT*REPRESENTATION*TIMING*SESSION-ORDER interaction (F_4, 68_ = 0.32, p = .863, partial η^2^ = .018). Bayes Factors for these interactions ranged between 3.08 and 30.09, providing strong to decisive evidence for a lack of effect of the session order on the effect of context on motor excitability.

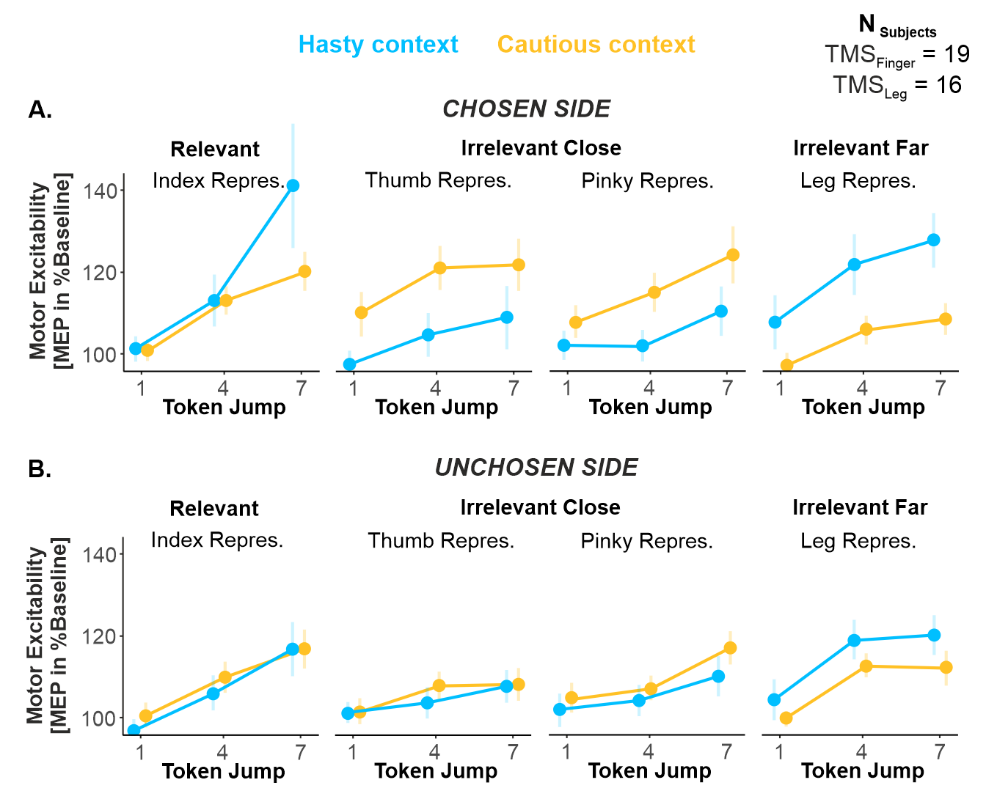

**S9 Figure (related to Figure 4): The effects of context were still present when exploiting the full, RT-unmatched dataset.** The RT-matching procedure described in S4 Figure ensured similar reaction times between the two contexts, but it raises a potential confound by emphasizing the slowest trials from the hasty context and the fastest trials from the cautious context. However, concerns about that confound are reduced by the observation that the same analyses performed on the full set of trials, without RT-matching, produced the same results. All individual and group-averaged numerical data exploited for S9 Figure are freely available at this link <https://osf.io/tbw7h/> (‘Figure_S9_Data.xlsx’).

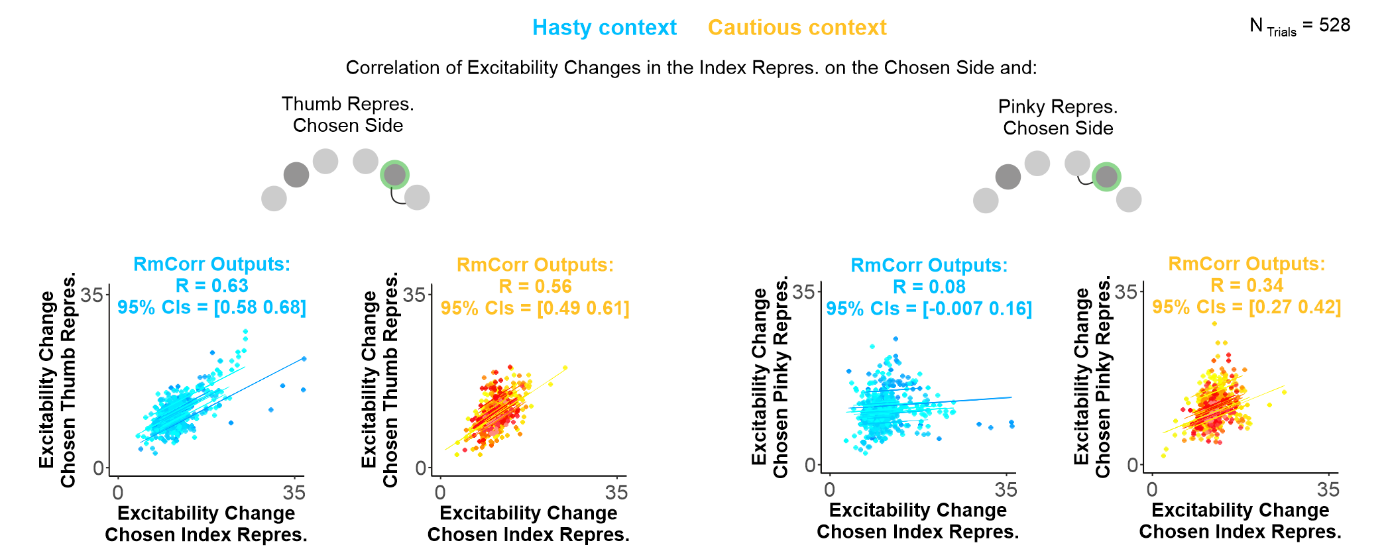

**S10 Figure (related to Figure 6): Single-trial correlations.** Example of four correlations obtained from the single-trial analysis. We pooled the trials of all subjects together (normalized to baseline; N_Subjects_ = 16), providing us with a large pool of data points (N_Trials_ = 528) and applied a repeated-measures correlation (rmCorr) analysis. RmCorr accounts for non-independence among observations using analysis of covariance (ANCOVA) to statistically adjust for inter-individual variability. By removing measured variance between-participants, rmCorr provides the best linear fit for each participant using parallel regression lines (the same slope) with varying intercepts, as can be seen in each cloud of points. As indicated in the methods section, normalized single-trial data were squared root-transformed to enhance the normality of the distributions (although similar findings were obtained on non-transformed data). As evident on this figure, excitability changes in the chosen index representation positively co-varied with changes in other finger representations (here the thumb and pinky representations of the chosen side). Further, while the strength of the correlation between the index and the thumb representation was comparable in the hasty and cautious contexts (R = 0.63 and 0.56, 95 % CIs = [0.58 0.68] and [0.49 0.61], respectively), the correlation between the index and the pinky representation was significantly weaker in the former than the latter context (R = 0.08 and R = 0.34, 95 % CIs = [-0.007 016] and [0.27 0.42], respectively). A similar decorrelation was observed between the chosen index and the index and thumb representations of the unchosen side (see Figure 6 in the main text). All individual and group-averaged numerical data exploited for S10 Figure are freely available at this link <https://osf.io/tbw7h/> (‘Figure_6&S10_Data.xlsx’).

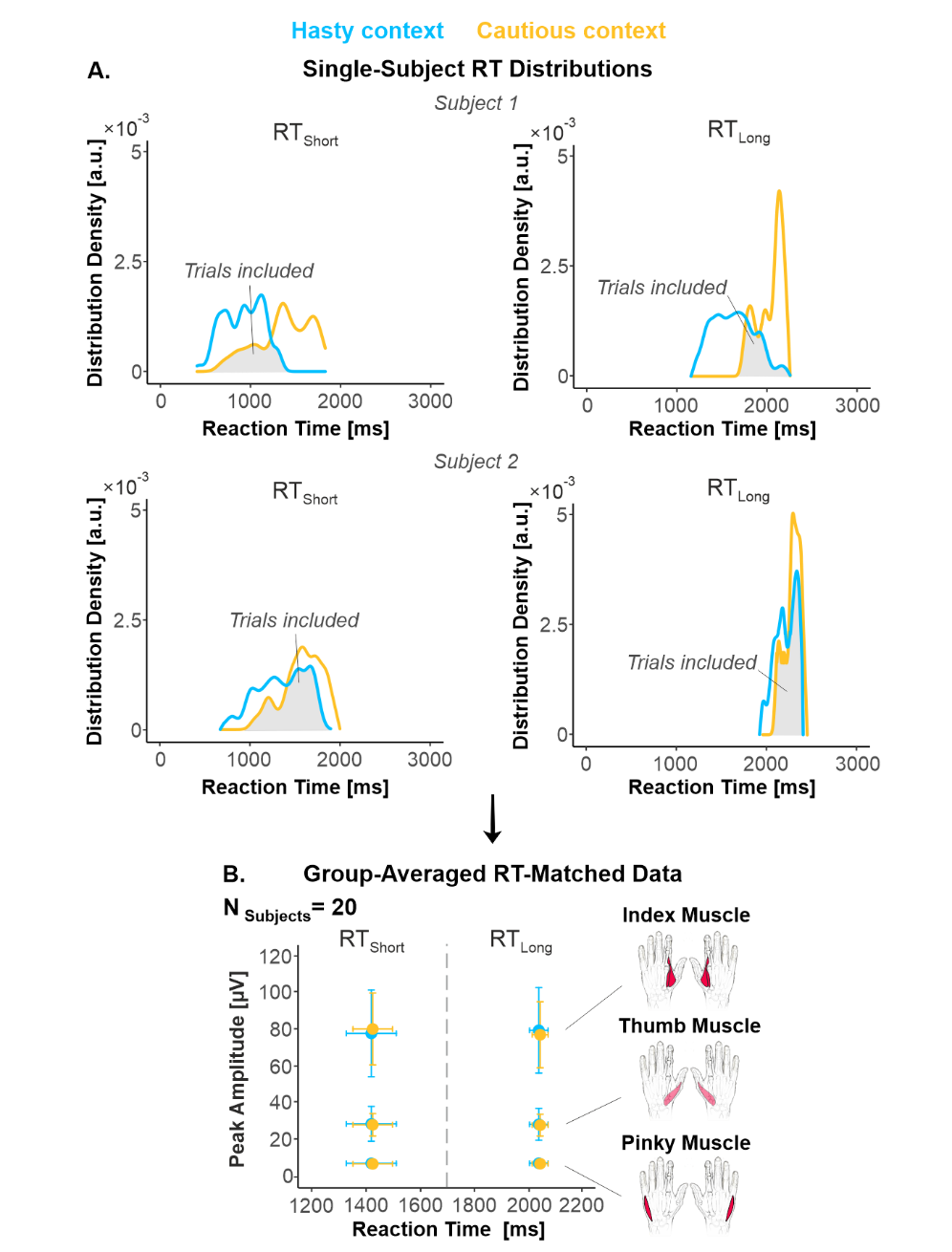

**S11 Figure: RT-median split and RT-matching procedures on movement vigor data.** **A. Distributions of included trials.** In order to investigate the effect of elapsed time on this EMG peak amplitude in each context and in every subject, we split the trials into two subsets according to whether they were associated with short or long RTs, using a median-split procedure (RT_Short_ and RT_Long_ trials, left and right panels, respectively). Further, to prevent EMG peak amplitude from being affected by the difference in decision speed between each context, we homogenized the RT distributions across contexts through a RT-matching procedure (same as described in S4 Figure on MEP data), both for RT_Short_ and RT_Long_ trials. **B. Group-averaged EMG peak amplitude.** Following the RT-matching procedure, we had to exclude 1 out of the 21 TMS_Finger_ subjects, as she/he ended up with no trial in a specific condition. As evident on the data averaged across the remaining 20 subjects, the included trials involved RTs that were comparable across contexts; this was true both for RT_Short_ (1414 ± 91 and 1422 ± 72 ms in hasty and cautious contexts, respectively) and for RT_Long_ trials (2030 ± 34 and 2034 ± 29 ms in hasty and cautious contexts, respectively). The figure also indicates the strong similarity of EMG peak amplitude values for RT_Short_ and RT_Long_ trials and for the hasty and cautious contexts. All individual and group-averaged numerical data exploited for S11 Figure are freely available at this link <https://osf.io/tbw7h/> (‘Figure_S11_Data.xlsx’).
